# Genotypic diversity, circulation patterns, and co-detections among rhinoviruses in Queensland, 2001

**DOI:** 10.1101/334334

**Authors:** Katherine E. Arden, Ristan M. Greer, Claire Y.T. Wang, Ian M. Mackay

**Affiliations:** Child Health Research Centre, The University of Queensland, Brisbane, Queensland, Australia; Faculty of Medicine, The University of Queensland, Brisbane, Queensland, Australia; Centre for Children’s Health Research, Children’s Health Queensland South Brisbane, Queensland, 4101, Australia

**Keywords:** Rhinovirus, RV, epidemiology, diagnostics, virus interactions, prevalence

## Abstract

Rhinoviruses (RVs) occur more frequently than other viruses and more often in people displaying symptoms than in those without. RVs exacerbate chronic airway disease and confound the clinical diagnosis of influenza-like illness. We sought to estimate the spectrum of RV diversity, RV species seasonality and to breakdown RV involvement in respiratory virus co-detections by comprehensive molecular testing of a convenience collection of airway sample extracts from patients with suspected respiratory infections, collected during 2001.

RVs were the most common virus detected. We were able to genotype ∼90% of RV detections, identifying 70 distinct RVs, spanning all three species. RV-Bs were under-represented. We found RV species co-circulated at times, although one species usually dominated. Each species displayed a bimodal distribution.

Notably, RVs and influenza A viruses (IFAV) seldom co-occurred, supporting their roles as primary pathogens of the airway among acutely ill infants. Whether RV circulation has a moderating or controlling effect on the IFAV season or is controlled by it cannot be determined from these data.

Despite the frequent perception that RVs commonly co-occur with another virus, our findings indicated this was not the case. Nearly 80% of RV detections occurred alone. Understanding more about population-level interference between viruses may allow us to harness aspects of it to generate a non-specific antiviral intervention that mimics a putative protective effect.

For routine respiratory virus screening to best serve the patient, RV testing should be a principal component of any acute respiratory illness testing algorithm throughout the year.

## Introduction

Rhinoviruses (RVs) are the largest related assemblage of genetically and antigenically distinct respiratory pathogens known, comprising 168 genotypes. These picornavirus species often occur in symptomatic young children from community and hospital populations where they create a sizable burden for management and are a frequent trigger of wheeze.^1-3^ Picornavirus infections can more often be observed among children presenting to an emergency department than other respiratory virus infections.^4^ Nearly three quarters of respiratory virus detection episodes during the first 28 days of life are due to RVs and over half are symptomatic.^3^ Until recently, much of what is known of RV diversity, epidemiology and clinical impact was determined using human studies and cell culture methods in the 1950s to 1980s. In 2006, the molecular discovery of a genetic clade of RVs that could only be cultured *in vitro* using sophisticated air-liquid interface cultures was subsequently ratified as a third species, *Rhinovirus C* (RV-C).^5, 6^ To date, RV-C has added 56 distinct RV types to the genus *Enterovirus*, family *Picornaviridae*. As more is learned, conclusions reached by some earlier studies have required re-examination and confirmation.^5-7^

We aimed to estimate the spectrum of RV genotypes, species seasonality and RV involvement in co-detections in Queensland using a convenience collection of airway sample extracts from patients with suspected respiratory infections, collected during 2001 and tested using molecular tools expected to account for all RV species.

## Methods

### Specimen extracts

Specimens extracts (n=1179; 94% nasopharyngeal aspirates) for retrospective RV testing originated from an unselected sample of mostly young patients (57.5% male) aged one day to 90.1 years (average 5.4 years, median 1.5 years) from across Queensland who presented to a hospital or clinic serviced by Pathology Queensland, Queensland Health (PQ), with symptoms of acute respiratory infection during 2001. There was no systematic sampling protocol used, thus our sample did not represent the total population of extracts received by PQ for testing in 2001. Extracts included nasopharyngeal aspirates (85.5%), lavage (9.5%), endotracheal aspirates (2.2%) and swabs (1.3%). Nucleic acids had been previously extracted and stored at -80°C as previously described.^8^ This study was approved by the Queensland Children’s Health Services human research ethics committee (#HREC/06/QCRH/097) and UQ medical research ethics committee #2009000039.

### Virus screening and RV species and type designation

Extracts had been previously tested by PQ using direct or culture-amplified direct fluorescent assay to detect respiratory syncytial virus (RSV), adenoviruses (AdV), parainfluenza viruses (PIVs) and influenza viruses A & B (IFAV, IFBV).^9^ PCR was not in routine use at PQ in 2001. This study applied additional previously described real-time RT-PCR (RT-rtPCR) assays to detect human metapneumovirus (MPV)^8, 10^ and human coronaviruses (HCoV) 229E, HKU1, NL63 and OC43.^11^ RVs were detected using a modified version of a previously described RT-rtPCR.^12, 13^ The RV RT-rtPCR can also detect some human enterovirus (EV) types.

RV/EV-positive extracts were genotyped using a conventional nested VP4/VP2 RT-PCR assay incorporating previously described primers that include part of the 5’ untranslated region (UTR), viral protein coding regions VP4 and partial VP2.^14-16^ first-round RT-PCR used OneStep RT-PCR (QIAGEN, Australia) or SensiFAST Probe No-ROX One-Step (Bioline, Australia) kits and subsequent second-round PCR used Bioline MyTaq HS DNA polymerase kit. In cases of failure to amplify a VP4/VP2 amplicon (∼540bp), part of the 5’ untranslated region (5’UTR; ∼400nt) was amplified.^16^Amplicons were sequenced (BigDye sequencing kit v3.1, Applied Biosystems Pty. Ltd.) and, after removal of the primer sequence (Geneious Pro v6)^17^, RV and EV sequences were submitted to GenBank (accession numbers KF499366-KF499501, KF688607-KF688723). Curated and detailed methods are available online.^18-20^

A virus variant was described as an “untypeable RV” when a clean sequence could not be obtained from either genotyping assay, despite the extracts being repeatedly positive by RT-rtPCR assay. RV genotype was determined by best match using the following algorithm: when a query sequence returned BLAST comparison with ≥97% sequence identity in the 5’UTR or ≥90% (for RV-B and RV-C) or ≥91% (for RV-A) in the VP4/VP2 region, with members assigned to a given type, our sequence was assigned as being a variant of that RV genotype.

### Statistical methods

We provide descriptive statistics for these data, using counts and proportions. Following methods we described previously,^21^ univariate analysis was used to screen the relationships between picornavirus groups (RV-A, RV-B, RV-C or EV), season and demographic variables such as age and sex. For each virus, the probability of co-detection with another virus was assessed with Fisher’s exact test using 2 × 2 contingency tables. We used a threshold p value of <0.05 for a statistically significant association. Logistic regression was used to estimate any effect of age, sex, season and other circulating virus types on detection of RV.

## Results

### Virus detections

In 615 extracts (52.2% of those tested) at least one virus was detected. The most frequently detected virus or virus group was a picornavirus (Figure 1 and Table 1; suspected RVs or EVs; n=296; 25.1% of all extracts, 48.1% of all virus positives) followed by RSV (n=101; 8.6% of all extracts), MPV (n=85; 7.2%), IFAV (n=73; 6.2%), PIV (n=66, 5.6%; 54 were PIV-3), AdV (n=37; 3.1%), HCoV-OC43 (n=26; 2.2%), HCoV-HKU1 (n=5; 0.4%), HCoV-NL63 (n=4; 0.3%), or HCoV-229E (n=1; 0.1%). No IFBV was detected and we did not seek human bocaviruses, influenza C virus or PIV-4.

**Table 1.**
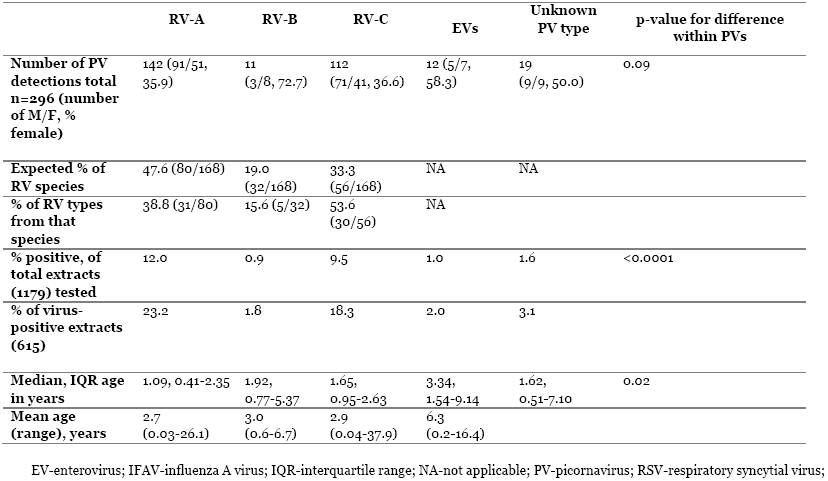
Features of picornavirus positive extracts.

**Figure 1.**
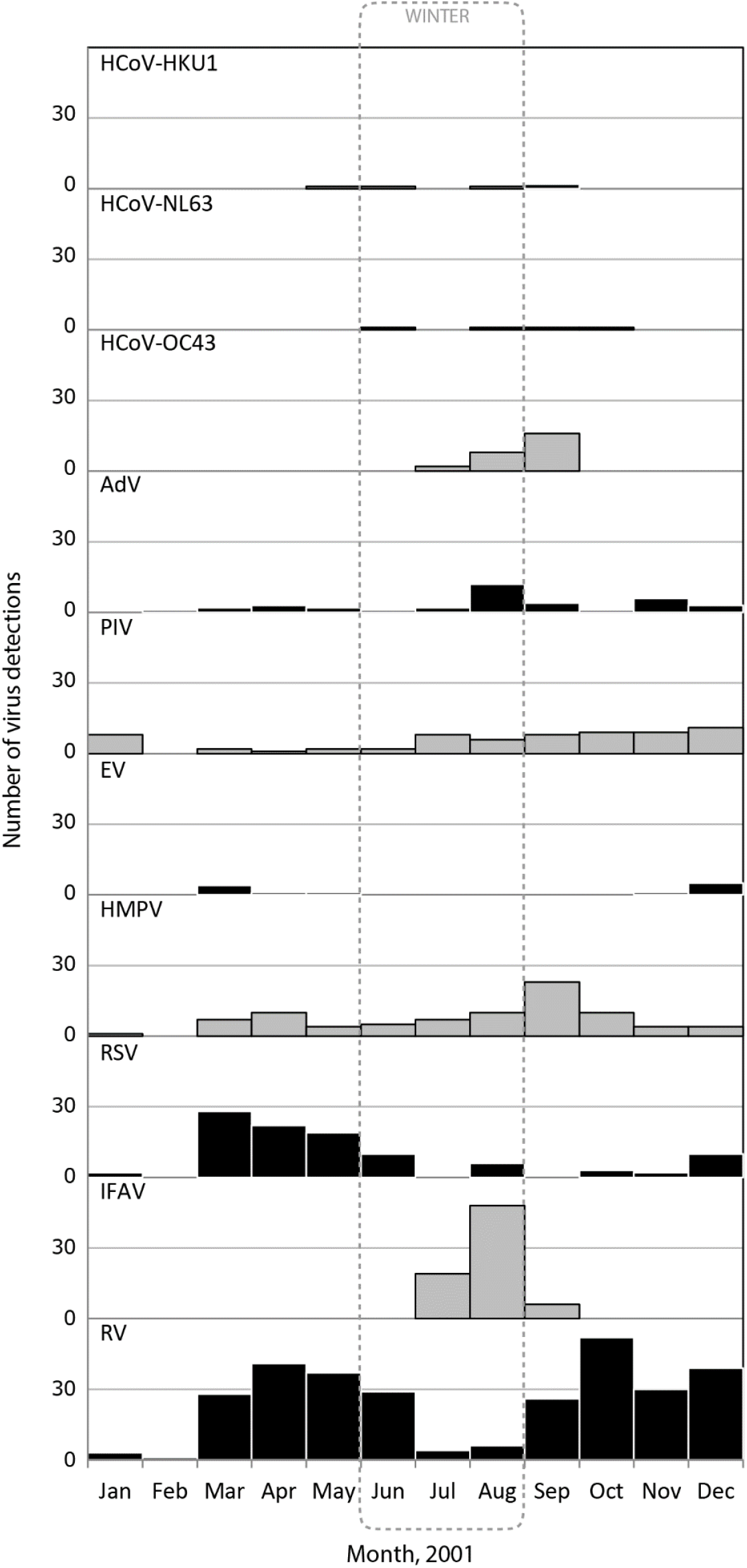
Number of extracts with a PCR-positive virus identification according to the month of sample collection and virus, 2001. Winter is defined by a dashed outline (June, July, August). HCoV-human coronavirus; AdV-any adenovirus; PIV-any parainfluenza virus; EV–any enterovirus; HMPV-human metapneumovirus; RSV-respiratory syncytial virus; IFAV-influenza A virus; RV-any rhinovirus. DOI 10.6084/m9.figshare.5549350.

### Demographic features of virus positives

The virus-positive population had a median age of 1.5 years (IQR 0.06-3.5 years); this was lowest for RV and RSV cases (2.9 and 2.4 years respectively). The age range varied among viruses in our population (Supplementary Figure 1). Fewer than 5% of neonates or adults were RV positive (Supplementary Figure 2).

There are more females in the Australian population (0.9:1.0^22^), but more males in the total patient sample (678/1179, 57.5%, p<0.0001 for expected proportion of 50%). In the context of total patient extracts, 180/678 (26.6%) extracts tested from males were picornavirus positive compared with 116/501 (23.0%) of extracts from females (p=0.20). Similar results were seen in the context of the 615 virus-positive extracts. Of the 296 picornavirus positive extracts, 60.8% were from males and 39.2% from females, but this difference in proportion was comparable to the sex proportions in the picornavirus negative group and was not significant (p=0.32).

### RV speciation and genotyping

An RV or EV type was assigned to 278 (93.9%) of 296 extracts (25.1%) that were picornavirus-positive by our screening RT-rtPCR. Most picornaviruses were RV-As and RV-Cs (Table 1). In total, 70 distinct RV types circulated at our site, including variants of 4 types that could not be assigned to an existing genotype (Supplementary Table 1). 12 genotypes represented 50.7% of those found and 3 genotypes belonged to the “minor group” of receptor-classified RVs. There are 32 known RV-B types therefore 19.2% of RV detections were expected if RVs are distributed randomly. However, only 3.7% of picornaviruses were RV-Bs, which is a significant under-representation (p<0.001). To ensure both genotyping methods performed as expected, we determined both 5’UTR and VP4/VP2 sequences for 10 RV positives and typing assignments agreed between targets using our thresholds.

### Seasonality

Total virus detections did not present with any clear seasonality but individual viruses and virus groups did. Most detections were in spring and autumn (61.2% of positives; Figure 1, Figure 2). RSV detections peaked in autumn, PIVs, HMPV and HCoV-OC43 in spring, AdV and IFAV during late winter and EVs during summer and autumn.

**Figure 2.**
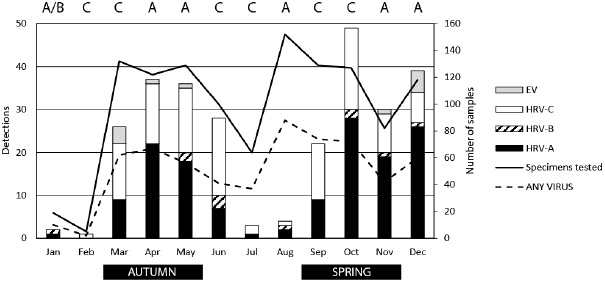
Number of extracts and virus detections. The total number of extracts tested (solid line, right y-axis) and the number of any virus detections (broken line, right y-axis) as well as the number of each RV species or EV detected (bar graph, left y-axis), per month, 2001. The predominating RV species is described at the top and key seasons along the bottom. DOI; 10.6084/m9.figshare.6388850.

Inspection of epidemic curves suggested some relationships between viral seasons. Spearman correlations supported these impressions, showing that both RV and RSV seasons were negatively associated with IFAV season (Supplementary Table 2). Positive correlations included RV with RSV; EV with RSV; AdV with PIVs; HMPV with IFAV, HCoV-OC43, HCoV-NL63 and PIV

RVs were detected in all seasons, but a bimodal distribution was evident with peaks in spring (36.5% of all picornaviruses) and autumn (35.8%) and a trough in winter (13.2%; summer extract numbers were low, so prevalence was difficult to determine; Figure 2). We observed seasonal exchange between RV species; RV-As and RV-Cs swapped dominance throughout the year. Although their numbers were lower, RV-Bs (none detected in 4/12 months) and EVs (none detected in 7/12 months) exchanged peak positions of prevalence throughout the year.

### Analysis of co-detections

Most viruses occurred as single detections (n=539; 87.6% of virus positives; Figure 3). Co-detections were found in 76 extracts, 71 with 2 viruses detected and 5 with 3 viruses detected; no extract had more than 3 viruses identified. Viral co-detection totals also exhibited a bimodal peak (Supplementary Figure 3).

**Figure 3.**
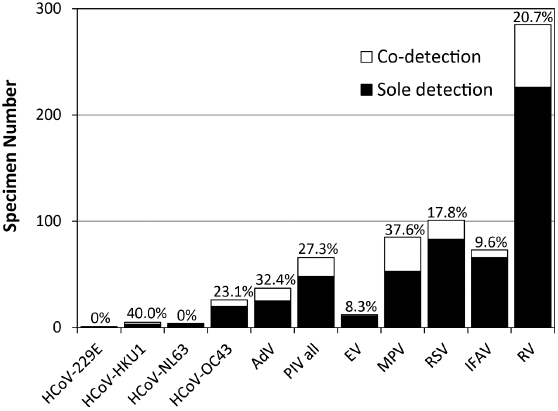
The number of single (filled bars) and co-detections (open bars) for each respiratory virus identified during 2001. The proportion of each virus’s detections that occur with another virus is highlighted. DOI 10.6084/m9.figshare.6390971.

Associations of RVs with other viruses, sex and season, were examined using logistic regression models, and results were consistent with those from the univariate analyses (Supplementary Table 3). Most RV detections occurred as single detections (n=225; 78.9% of RVs), most often in autumn (when RSV detections also peaked) and spring (when MPV and PIV detections peaked) (Figure 1).

Detection of more than one virus occurred in 76 extracts. RVs were the most frequently detected viruses overall and were involved in the greatest total number of co-detections (n=60; 20.3% of all 296 RV positive extracts) (Figure 3; Supplementary Table 4). Most (n=44; 73.3%) RV co-detections occurred in those 2-years of age or younger, and none of the 60 extracts originated from adults (>14-years of age). The proportion of each RV species involved in co-detections differed (p<0.0001). Of the RV-As, 21 of 142 (14.8%) were co-detections while 7 of 11 (63.6%) RV-Bs and 22 of 112 (19.6%) RV-Cs were co-detections. One of the 12 EVs (8.3%) was a co-detection. RV-Bs were more likely to occur as co-detections than RV-As (p=0.001), RV-Cs (p=0.003) or EVs (p=0.009). Ten of the 19 (52.6%) untypeable picornaviruses were co-detections. Most RV co-detections were with paramyxoviruses (MPV, n=21; RSV, n=17; PIV, n=16) followed by AdV (n=5), IFAV (n=2) and HCoV-HKU1 (n=1). There were no co-detections identified between RV or RV and EV species (p<0.004 for all comparisons) except for RV-B and EV. Given a positive detection of EV, 0/11 (0%) extracts were positive for RV-B compared with 12/285 (4.2%) extracts negative for RV-B, p=1.0.

It was important to look at co-detections as a proportion of each virus, not just RVs. Other viruses were involved in a similar or greater proportion of co-detections compared to the RVs (Figure 3. IFAV had one of the lowest proportion of co-detections (9.6%) as did the EVs (8.3%) while HCoV-NL63 was not involved in any co-detections. RVs were significantly less likely than expected to be co-detected with IFAV (p<0.0001; Supplementary Table 4). PIVs were less likely to be detected with RSV (p=0.05) or IFAV (p=0.03), and RSV was less likely to be found with IFAV (p=0.002) than expected.

## Conclusions

Seventy distinct RV genotypes circulated among this symptomatic convenience population. Levels of genotypic diversity vary among other studies; some are higher than we observed, such as during the first year of life among the childhood origins of asthma cohort ^23^, but many observe fewer distinct genotypes.^24-26^ Studies which do not account for RV species or genotype risk overestimating the length of RV shedding, falsely assuming persistence rather than sequential infection.^27, 28^ Among our sample, RVs dominated in prevalence compared to all other respiratory viruses across most of 2001, notably waning during the seasonal IFAV epidemic.

Most picornavirus positive extracts were genotyped, a fact that agreed with some studies, but not others. ^23, 29 28, 30-32^ Lower rates of genotyping success in community studies may be due to lower viral loads among less severely ill subjects. We propose that our combination of real-time RT-PCR screening and conventional nested-PCR genotyping methods are an effective starting point for genotyping ∼90% of RVs among ill populations, even from extracts stored for over a decade.

Others have found RV-Cs to be the dominant RV species in hospital-associated asthma exacerbations.^1, 33^ We have previously found them to be numerically dominant but RV-A to be more clinically significant in non-hospitalised asthmatics.^34^ We observed RV-B detections to be a significantly smaller proportion of total RV detections than was expected. We have previously noted this in studies of hospitalised populations.^29, 34^ It is not clear why RV-Bs are under-represented. Perhaps the RV-Bs are less severe pathogens ^23, 26^ or their infections contribute disease to a different population than that tested here. It is noteworthy that RV-Bs are also under-represented in community populations.^28, 35^

While the RV species often co-circulated during 2001, one species usually predominated each month, changing from month to month. Each species displayed bimodal peaks in autumn and spring and a trough in winter, which aligned with IFAV circulation. Exchange of prevailing species and turnover of prevalent genotypes is supported by other studies that employ molecular typing and span 12 months or more.^25 15, 36-39^ Shorter cross-sectional studies may misrepresent association between disease severity and RV species or genotype by failing to capture year-to-year variation in RV diversity.

Whether RV circulation has a moderating or controlling effect on the IFAV season or is controlled by it, cannot be determined from these data however these viruses seldom seasonally co-occurred. We and others have seen this and related interactions, termed virus interference, in other populations.^29, 40, 41^ Virus interference is a consistent feature that some have attributed to RV dominance.^42, 43^ It remains unclear whether this interference pattern is a feature associated with a mostly paediatric symptomatic population, a high proportion (86%) of nasopharyngeal extracts, whether influenza type or subtype plays a role or if pre-existing RV immunity can influence interference. Influenza may control a population’s susceptibility to other respiratory viruses, blocking their spread because of strong a population-level innate immune response generated by influenza strains during an annual epidemic “flu season”.

Initiation of seasonal influenza epidemics is mostly associated with rate of influenza virus change, strain replacement, availability of susceptible hosts and climatic conditions.^44, 45^ Further study of virus interference is warranted and could inform our understanding of human respiratory diseases that exhibit temporal trends such as exacerbations of asthma and chronic obstructive pulmonary disease, otitis media and allergic rhinitis. Understanding a mechanism for interference may allow us to harness it to generate a non-specific antiviral intervention that mimics this putative protection.

We found nearly five times more co-detections in our study than were reported from among the first viral detections of those enrolled in a community birth cohort from Brisbane.^3, 27^ EVs were observed to have an especially low proportion of co-detections in our 2001 sample. As reported in our earlier studies of populations from different years and Australian locations, we again observed that detection of an IFAV, EV, RV or RSV occurred with reduced likelihood of co-detection of any other virus in that patient’s sample.^21, 29^ There was a higher likelihood of co-detection among extracts positive for an AdV or MPV. Numbers were small for the other viruses. The accepted understanding of IFAV directly associated with severe illness was supported here by a low proportion of co-detections.

Infants from one to 12 months of age were most likely to be affected by RV or RSV in our convenience sample population; this agreed with findings from the Brisbane community cohort.^3^ RV-positive children were second youngest after RSV-positive children while IFAV-positive children were generally older than those positive for any other virus. Interestingly, it is often children younger than 2 years who are affected by severe influenza.^46^ Most RV detections (single or multiple) occurred in infants, the largest contributing population. It is unknown whether the viruses detected in this population reflect those that concurrently circulated among mild illness or subclinical community infections during 2001.

This analysis provides useful comparative baseline data for ongoing and future analyses.^28, 47^ There is no denominator from which to determine the rates of virus infection nor is there a specific clinical definition of each infected person available. Neither clinical impact nor admission status was sought in this study, but sampling was due to a clinical need. The small number of extracts available from January and February (summer) limited the strength of the RV analyses during this period. Also, the absence of testing for bocaviruses may have reduced the positive associations noted among the respiratory viruses because bocaviruses are frequent partners in extracts positive for more than one respiratory virus.^5, 21^

Despite the frequent perception that RVs commonly co-occur with another virus, our findings here and elsewhere indicate this is less likely to occur for RVs than for other respiratory viruses.^21, 29^ That nearly 80% of RV detections occurred in the absence of another virus once again supports the assertion that RVs are primary pathogens of the airway among acutely ill infants. If a respiratory tract sample is to be collected for any testing and if routine screening of respiratory viruses is to serve its patients best, RV should be a principal component of any acute respiratory illness testing algorithm throughout the year, with allowance for their relative replacement by IFAV cases during winter.

## Authorship

IMM planned the study, conducted experiments, analysed data, acquired funding and wrote the manuscript; KEA conducted experiments, analysed data and co-wrote the manuscript; CYTW conducted experiments, analysed data; RMG performed statistical analyses and co-write the manuscript. All authors reviewed, revised and approved the final version.

## Acknowledgements

We thank Pathology Queensland Central for providing clinical specimen nucleic acid extracts, Michael Witt for aid with data collection and Stephen Lambert for helpful discussions. Special thanks to Cassandra Faux for outstanding contributions to the laboratory work.

## References

1. Miller EK, Khuri-Bulos N, Williams JV, Shehabi AA, Faouri S, Al JI, et al. Human rhinovirus C associated with wheezing in hospitalised children in the Middle East. J Clin Virol. 2009;46:85–89.

2. Piotrowska Z, V zquez M, Shapiro ED, Weibel C, Ferguson D, Landry ML, et al. Rhinoviruses are a major cause of wheezing and hospitalization in children less than 2 years of age. Pediatric Infectious Disease Journal. 2009;28:25–29.

3. Sarna M, Ware RS, Lambert SB, Sloots TP, Nissen MD, Grimwood K. Timing of First Respiratory Virus Detections in Infants: A Community-Based Birth Cohort Study. J Infect Dis. 2018;217:418–27.

4. Lambert SB, Allen KM, Druce JD, Birch CJ, Mackay IM, Carlin JB, et al. Community epidemiology of human metapneumovirus, human coronavirus NL63, and other respiratory viruses in healthy preschool-aged children using parent-collected specimens. Pediatrics. 2007;120:e929–e37.

5. Arden KE, McErlean P, Nissen MD, Sloots TP, Mackay IM. Frequent detection of human rhinoviruses, paramyxoviruses, coronaviruses, and bocavirus during acute respiratory tract infections. Journal of medical virology. 2006;78:1232–40.

6. Simmonds P, McIntyre CL, Savolainen-Kopra C, Tapparel C, Mackay IM, Hovi T. Proposals for the classification of human rhinovirus species C into genotypically-assigned types. JGenVirol. 2010;91:2409–19.

7. Royston L, Tapparel C. Rhinoviruses and Respiratory Enteroviruses: Not as Simple as ABC. Viruses. 2016;8.

8. Mackay IM, Bialasiewicz S, Jacob KC, McQueen E, Arden KE, Nissen MD, et al. Genetic diversity of human metapneumovirus over 4 consecutive years in Australia. J Infect Dis. 2006;193.

9. Syrmis MW, Whiley DM, Thomas M, Mackay IM, Williamson J, Siebert DJ, et al. A sensitive, specific and cost-effective multiplex reverse-transcriptase-PCR assay for the detection of seven common respiratory viruses in respiratory samples. Journal of Molecular Diagnostics. 2004;6:125–31.

10. Maertzdorf J, Wang CK, Brown JB, Quinto JD, Chu M, de Graaf M, et al. Real-time reverse transcriptase PCR assay for detection of human metapneumoviruses from all known genetic lineages. Journal of Clinical Microbiology. 2004;42:981– 86.

11. Mackay IM, Arden KE, Speicher DJ, O’Neill NT, McErlean PK, Greer RM, et al. Co-circulation of four human coronaviruses (HCoVs) in Queensland children with acute respiratory tract illnesses in 2004. Viruses. 2012;4:637–53.

12. Lu X, Holloway B, Dare RK, Kuypers J, Yagi S, Williams JV, et al. Real-time reverse transcription-PCR assay for comprehensive detection of human rhinoviruses. Journal of Clinical Microbiology. 2008;46:533–39.

13. Arden KE, Mackay IM. Newly identified human rhinoviruses: molecular methods heat up the cold viruses. Rev Med Virol. 2010;20:156–76.

14. Savolainen C, Blomqvist S, Mulders MN, Hovi T. Genetic clustering of all 102 human rhinovirus prototype strains: serotype 87 is close to human enterovirus 70. Journal of General Virology. 2002;83:333–40.

15. Wisdom A, McWilliam Leitch EC, Gaunt E, Harvala H, Simmonds P. Screening and comprehensive VP4/2-typing of human rhinoviruses (HRVs) and Enteroviruses: Comprehensive VP4-VP2 typing reveals high incidence and genetic diversity HRV Species C. Journal of Clinical Microbiology. 2009;47:3958–67.

16. Gama RE, Horsnell PR, Hughes PJ, North C, Bruce CB, Al-Nakib W, et al. Amplification of rhinovirus specific nucleic acids from clinical samples using the polymerase chain reaction. Journal of medical virology. 1989;28:73-77.

17. Geneious.

18. Mackay IM. Human rhinovirus screening conventional RT-PCR (“Gama assay”). protocols.io2018.

19. Mackay IM, Wang CYT, Arden KE. Respiratory picornavirus genotyping conventional nested RT-PCR (“Wisdom VP42 assay”). protocols.io2018.

20. Mackay IM. Human rhinovirus screening real-time RT-PCR (“modified Lu assay”). protocols.io2018

21. Greer RM, McErlean P, Arden KE, Faux CE, Nitsche A, Lambert SB, et al. Do rhinoviruses reduce the probability of viral co-detection during acute respiratory tract infections? JClinVirol. 2009;45:10–15.

22. Australian Bureau of S. Population by Age and Sex, Australia, States and Territories. Australian Bureau of Statistics; 2013.

23. Lee WM, Lemanske RF, Jr., Evans MD, Vang F, Pappas T, Gangnon R, et al. Human rhinovirus species and season of infection determine illness severity. Am J Respir CritCare Med. 2012;186:886–91.

24. Marcone DN, Culasso A, Carballal G, Campos R, Echavarria M. Genetic diversity and clinical impact of human rhinoviruses in hospitalized and outpatient children with acute respiratory infection, Argentina. J Clin Virol. 2014;61:558–64.

25. Stelzer-Braid S, Tovey ER, Willenborg CM, Toelle BG, Ampon R, Garden FL, et al. Absence of back to school peaks in human rhinovirus detections and respiratory symptoms in a cohort of children with asthma. Journal of medical virology. 2016;88:578–87.

26. Tran DN, Trinh QD, Pham NT, Pham TM, Ha MT, Nguyen TQ, et al. Human rhinovirus infections in hospitalized children: clinical, epidemiological and virological features. Epidemiol Infect. 2016;144:346–54.

27. Byington CL, Ampofo K, Stockmann C, Adler FR, Herbener A, Miller T, et al. Community Surveillance of Respiratory Viruses Among Families in the Utah Better Identification of Germs-Longitudinal Viral Epidemiology (BIG-LoVE) Study. Clin Infect Dis. 2015;61:1217–24.

28. Sarna M, Alsaleh A, Lambert SB, Ware RS, Mhango LP, Mackay IM, et al. Respiratory Viruses in Neonates: A Prospective, Community-Based Birth Cohort Study. The Pediatric Infectious Disease Journal. 2016;Publish Ahead of Print.

29. Mackay IM, Lambert SB, Faux CE, Arden KE, Nissen MD, Sloots TP, et al. Community-wide, contemporaneous circulation of a broad spectrum of human rhinoviruses in healthy Australian preschool-aged children during a 12-month period. J Infect Dis. 2012;207:1433–41.

30. Smuts HE, Workman LJ, Zar HJ. Human rhinovirus infection in young African children with acute wheezing. BMCInfectDis. 2011;11:65.

31. Miller EK, Williams JV, Gebretsadik T, Carroll KN, Dupont WD, Mohamed YA, et al. Host and viral factors associated with severity of human rhinovirus-associated infant respiratory tract illness. J AllergyClin Immunol. 2011;127:883–91.

32. Principi N, Zampiero A, Gambino M, Scala A, Senatore L, Lelii M, et al. Prospective evaluation of rhinovirus infection in healthy young children. J Clin Virol. 2015;66:83–9.

33. Miller EK, Edwards KM, Weinberg GA, Iwane MK, Griffin MR, Hall CB, et al. A novel group of rhinoviruses is associated with asthma hospitalizations. Journal of Allergy and Clinical Immunology. 2009;123:98–104.

34. Arden KE, Chang AB, Lambert SB, Nissen MD, Sloots TP, Mackay IM. Newly Identified Respiratory Viruses in Children with Non-hospitalised Asthma Exacerbation. Journal of medical virology. 2010;82:1458–61.

35. Muller L, Mack I, Tapparel C, Kaiser L, Alves MP, Kieninger E, et al. Human Rhinovirus Types and Association with Respiratory Symptoms During the First Year of Life. Pediatr Infect Dis J. 2015;34:907–9.

36. Xiang Z, Gonzalez R, Wang Z, Xiao Y, Chen L, Li T, et al. Human rhinoviruses in Chinese adults with acute respiratory tract infection. JInfect. 2010;61:289–98.

37. Xiang Z, Gonzalez R, Xie Z, Xiao Y, Liu J, Chen L, et al. Human rhinovirus C infections mirror those of human rhinovirus A in children with community-acquired pneumonia. JClinVirol. 2010;49:94–99.

38. Lau SKP, Yip CC, Lin AW, Lee RA, So LY, Lau YL, et al. Clinical and molecular epidemiology of human rhinovirus C in children and adults in Hong Kong reveals a possible distinct human rhinovirus C subgroup. J Infect Dis. 2009;200:1096– 103.

39. van der Zalm MM, Wilbrink B, van Ewijk BE, Overduin P, Wolfs TF, van der Ent CK. Highly frequent infections with human rhinovirus in healthy young children: a longitudinal cohort study. J Clin Virol. 2011;52:317–20.

40. Yang Y, Wang Z, Ren L, Wang W, Vernet G, Paranhos-Baccala G, et al. Influenza A/H1N1 2009 pandemic and respiratory virus infections, Beijing, 2009-2010. PLoSOne. 2012;7:e45807.

41. Ånestad G, Vainio K, Hungnes O. Interference between outbreaks of epidemic viruses. Scandinavian Journal of Infectious Diseases. 2007;39:653–54.

42. Casalegno JS, Ottmann M, Duchamp MB, Escuret V, Billaud G, Frobert E, et al. Rhinoviruses delayed the circulation of the pandemic influenza A (H1N1) 2009 virus in France. ClinMicrobiolInfect. 2010;16:326–29.

43. Linde A, Rotzen-Ostlund M, Zweygberg-Wirgart B, Rubinova S, Brytting M. Does viral interference affect spread of influenza? EuroSurveill. 2009;14.

44. Furuse Y, Oshitani H. Mechanisms of replacement of circulating viruses by seasonal and pandemic influenza A viruses. Int J Infect Dis. 2016;51:6–14.

45. Bedford T, Riley S, Barr IG, Broor S, Chadha M, Cox NJ, et al. Global circulation patterns of seasonal influenza viruses vary with antigenic drift. Nature. 2015;523:217–20.

46. Buchan SA, Chung H, Campitelli MA, Crowcroft NS, Gubbay JB, Karnauchow T, et al. Vaccine effectiveness against laboratory-confirmed influenza hospitalizations among young children during the 2010-11 to 2013-14 influenza seasons in Ontario, Canada. PLoS One. 2017;12:e0187834.

47. Lambert SB, Ware RS, Cook AL, Maguire FA, Whiley DM, Bialasiewicz S, et al. Observational Research in Childhood Infectious Diseases (ORChID): a dynamic birth cohort study. BMJOpen. 2012;2.

